# Analysis of 19 Highly Conserved *Vibrio cholerae* Bacteriophages Isolated from Environmental and Patient Sources Over a Twelve-Year Period

**DOI:** 10.1101/313346

**Authors:** Angus Angermeyer, Moon Moon Das, Durg Vijai Singh, Kimberley D. Seed

## Abstract

The *Vibrio cholerae* biotype ‘El Tor’ is responsible for all current epidemic and endemic cholera outbreaks worldwide. These outbreaks are clonal and are hypothesized to originate from the coastal areas near the Bay of Bengal where the lytic bacteriophage ICP1 specifically preys upon these pathogenic outbreak strains. ICP1 has also been the dominant bacteriophage found in cholera patient stool since 2001. However, little is known about its genomic differences between ICP1 strains collected over time. Here we elucidate the pan-genome and phylogeny of ICP1 strains by aligning, annotating and analyzing the genomes of 19 distinct isolates collected between 2001 and 2012. Our results reveal that ICP1 isolates are highly conserved and possess a large core-genome as well as a smaller, somewhat flexible accessory-genome. Despite its overall conservation, ICP1 strains have managed to acquire a number of unknown genes as well as a CRISPR-Cas system, which is known to be critical for its ongoing struggle for co-evolutionary dominance over its host. This study describes a foundation on which to construct future molecular and bioinformatic studies of this *V. cholerae*-associated bacteriophages.

## 1. Introduction

*Vibrio cholerae* is a globally-distributed bacterium and the causal agent of the disease cholera, a potentially severe intestinal illness that affects ~1-5 million people resulting in up to ~140,000 deaths annually [1]. The current (seventh) pandemic is comprised of *V. cholerae* biotype ‘El Tor’ (predominantly serotype O1) and is responsible for current endemic and epidemic disease [2]. Epidemic disease outbreaks sweep the globe periodically and have been traced back to a single lineage that has emerged from the Bay of Bengal region in multiple waves over the last half-century [3]. Despite the overall genetic heterogeneity of this lineage, in which individual outbreaks are nearly always clonal [4], there is an abundance of subtle variation and horizontal transfer that has been observed between outbreaks over time [5,6]. It is hypothesized that the Bay of Bengal serves as a reservoir where El Tor strains circulate throughout the year exchanging genetic material and undergoing ecological selection before infiltrating coastal communities. They are subsequently transported by infected individuals to larger cities where they can be transmitted globally [5,7]. This mechanism is thought to create a bottleneck for strains and result in the clonality of outbreaks.

Bacteriophage, viruses that uniquely infect bacteria, are extremely abundant in the environment where they can outnumber their prokaryotic hosts by several orders of magnitude [8]. As such, bacteriophage play a key role in the evolution of their hosts through both selection and phage-mediated lateral gene transfer [9]. These processes are likely to be very important to *V. cholerae* strain evolution in the Bay of Bengal as well. Previous work has identified a *V. cholera* O1-specific [10] lytic *myoviridae* bacteriophage (ICP1) to be of particular interest in this system [11]. In Bangladesh, ICP1 has been found in water samples [12,13] and it has been identified as the dominant phage in cholera patient stool samples since 2001 [11]. The persistence of this phage over time indicates that *V. cholerae* has strategies to limit ICP1 predation, and that ICP1 can evolve to overcome such defenses. Indeed, from this natural genetic laboratory, several complex and surprising adaptations/acquisitions have occurred in the race for survival between *V. cholerae* and ICP1. These include self-mobilizing chromosomal islands that can provide a rapid and efficient response to ICP1 infection [14] and the first known example of a bacteriophage-encoded CRISPR-Cas system [15]. Initial characterization of eight ICP1 isolates collected between 2001-2011 noted the relative low level of diversity and lack of major genomic rearrangements, deletions or insertions [11] (with the exception of its remarkable CRISPR-Cas acquisition [15]). Here we build upon the initial characterization of ICP1 to perform a comparative genomic analysis on 19 individual ICP1 isolates to reconstruct their phylogenetic relationships over time, identify the core and accessory genomes, and infer possible gene function where possible.

*V. cholerae* is an organism that affects millions and appropriately, it is well-studied with modern bioinformatic and sequencing tools. It is important that their concomitant bacteriophage are similarly studied to help us better elucidate the important role that bacteriophage likely play in the evolution of *V. cholerae* and the epidemiology of the ongoing cholera pandemic.

## 2. Materials and Methods

Nineteen ICP1 bacteriophage genomes were acquired from various sources (Table 1). Those genomes not specifically described below were downloaded from the NCBI GenBank database. Their metadata (if available) were used to inform our reporting of isolation source and isolation year. In cases where this information differed from what was reported in a genome’s original publication we deferred to the GenBank database. Isolation year was used to standardize the ICP1 bacteriophage naming convention: ICP1_YEAR_X where “X” is sequentially assigned letter (A-Z) based on the order in which genomes were named. The only exception is the original, ‘ancestral’ ICP1 isolate, which is simply referred to as ‘ICP1’ [11].

**Table 1.** ICP1 bacteriophage strains.

| Standardized Name | Previous Name | Isolation Year | Isolation Source | Genome Size (bp) | GenBank Accession | Genome Citation |
| --- | --- | --- | --- | --- | --- | --- |
| ICP1 | - | 2001 | Stool | 125,956 | HQ641347 | Seed *et al.* 2011 |
| ICP1_2001_A | - | 2001 | Stool | 124,826 | HQ641353 | Seed *et al.* 2011 |
| ICP1_2001_B | JSF1 | 2001 | Water | 126,082 | KY883636 | Naser *et al.* 2017 |
| ICP1_2001_C | JSF2 | 2001 | Water | 126,082 | KY883637 | Naser *et al.* 2017 |
| ICP1_2001_D | JSF4 | 2001* | Water* | 124,261 | KY065147 | Naser *et al.* 2017 |
| ICP1_2001_E | JSF5 | 2001* | Water | 132,142 | KY883634 | Naser *et al.* 2017 |
| ICP1_2001_F | JSF6 | 2001* | Water | 133,685 | KY883635 | Naser *et al.* 2017 |
| ICP1_2004_A | - | 2004 | Stool | 128,083 | HQ641354 | Seed *et al.* 2011 |
| ICP1_2005_A | - | 2005 | Stool | 129,373 | HQ641352 | Seed *et al.* 2011 |
| ICP1_2006_A | - | 2006 | Stool | 123,104 | HQ641351 | Seed *et al.* 2011 |
| ICP1_2006_B | - | 2006 | Stool | 123,097 | HQ641350 | Seed *et al.* 2011 |
| ICP1_2006_C | - | 2006 | Stool | 124,497 | HQ641349 | Seed *et al.* 2011 |
| ICP1_2006_D | - | 2006 | Stool | 124,497 | HQ641348 | Seed *et al.* 2011 |
| ICP1_2006_E | - | 2006 | Stool | 128,298 | TBD | This study |
| ICP1_2009_A | JSF13 | 2009 | Water | 128,814 | KY883638 | Naser *et al.* 2017 |
| ICP1_2011_A | - | 2011 | Stool | 126,861 | TBD | This study |
| ICP1_2011_B | - | 2011 | Stool | 125,128 | TBD | This study |
| ICP1_2011_C | JSF14 | 2011 | Water | 125,096 | KY883639 | Naser *et al.* 2017 |
| ICP1_2012_A | - | 2012 | Stool | 121,418 | TBD | This study |

\* GenBank metadata was reported in cases where it differed from the original publication.

Genomes not already available on GenBank include: ICP1_2006_E and ICP1_2011_A, which were isolated and sequenced as described previously [15]. ICP1_2011_A was assembled using CLC Genomics Workbench v10 (Qiagen, Redwood City, CA) and ICP1_2006_E was assembled with IDBA-UD v1.1.3 [16] (default settings) after fastq read filtering with USEARCH v10.0.240 [17] fastq_filter (-fastq_maxee_rate 0.001-fastq_maxns 1-fastq_truncqual 15-fastq_maxee 0.25). ICP1_2011_B was assembled from an existing diarrheal stool metagenomic sample [18] (SRA: PRJEB9150; Run: ERR866578) using USEARCH filtering and IDBA-UD *de novo* assembly as above (same parameters) to generate an incomplete ICP1 contig. That contig was then used as a reference sequence to reassemble the same filtered reads using IDBA-Hybrid [16] with default settings. Finally, ICP1_2012_A was isolated from a cholera patient stool sample collected in Silvassa, India [19]. Genomic DNA libraries were sequenced using an Illumina Mi-Seq (Genotypic Technology, Bangalore, India). Genome assembly was performed with CLC Genomics Workbench v10.

Whole-genome alignment was performed on all 19 genomes using progressiveMauve [20] (build: Feb 25 2015) with default settings. The Mauve xmfa alignment output file was converted to Phylip format using BioPython v1.71 [21] and a maximum likelihood, unrooted, phylogenetic tree was constructed using PhyML v20120412 [22] while calculating bootstrap support (-s BEST--rand_start--n_rand_starts 10-b 100). A companion phylogeny based on the concatenated core-genome (described below) was constructed using the same methods. Dendroscope3 v3.5.9 [23] was used to generate a tanglegram joining both trees with ICP1 ancestral set as the outgroup. The whole-genome alignment was also used to determine a consensus ICP1 sequence using CLC Genomics Workbench v10, which all ICP1 genomes were visually mapped back onto using BRIG v0.95 [24].

All extant CDS annotations, hereafter referred to as ‘Open Reading Frames’ (ORFs), in the ICP1 ancestral GenBank file were blasted (BLASTn v2.6.0 [25]) against every other ICP1 genomic sequence to find homologous ORFs. Putative hits were considered homologs, and annotated as such, if the following conditions were met: the subject hit was a complete ORF (start and stop codons; codon table=11), e-value≤ 1e-10, %identity≥85, and the two matched ORFs were within 10% sequence length of each other. After identifying existing homologs, *de novo* ORF prediction was performed on all genomes to find additional possible coding sequences using Prodigal v2.6.3 [26] with default settings and a confidence cutoff ≥95%. All newly identified putative-ORFs from each genome were then blasted against each other with the same conditions as above, grouped by homology and given an iterative numerical identifier based on their locations in the genome relative to the extant ORFs (i.e. ORF1, ORF2, ORF2.1, ORF2.2, ORF3, etc.). ORFs were then categorized into groups based on how many of the 19 genomes they were found it. If an ORF was present in all 19 genomes, it was considered part of the core-genome, otherwise, it was considered part of the accessory-genome. Core and accessory ORFs were also mapped to the BRIG alignment. The core-genome ORF protein sequences were concatenated by strain to create a core-genomic, pseudo-genome for each ICP1 strain and subjected to the phylogenetic analysis as above. The core and accessory ORFs were also queried against the NCBI’s Conserved Domain Database (https://www.ncbi.nlm.nih.gov/cdd
) to determine if any possessed interesting domain homology that wasn’t detected by BLAST alone.

## 3. Results

### 3.1. Genome Characteristics and Phylogeny

ICP1 genomes were analyzed from isolates collected over a 12-year period, from 2001 to 2012 (Table 1). The isolates were derived from both environmental water samples and patient stool samples collected in Bangladesh (n=18) and India (ICP1_2012_A). Genome length was slightly variable with an average of 126,384bp (stdev: 3008bp). The maximum likelihood phylogenetic analysis grouped both the whole-genome and core-genome alignments into several general clusters (Figure 1). Cluster A contains the five most recently isolated strains as well as ICP1_2004_A. This cluster also contains five of the seven total CRISPR-positive strains in the dataset. Cluster B contains four of the six 2001 isolates, ICP1_2006_E and two of the CRISPR-positive strains. Clusters C and D contain isolates from intermediate years 2005 and 2006, with one CRISPR-positive represented in cluster C. The CRISPR-positive strain ICP1_2001_F did not cluster with other isolates. ICP1 ancestral and ICP1_2001C also cluster closely together but are not specifically highlighted.

**Figure 1.**
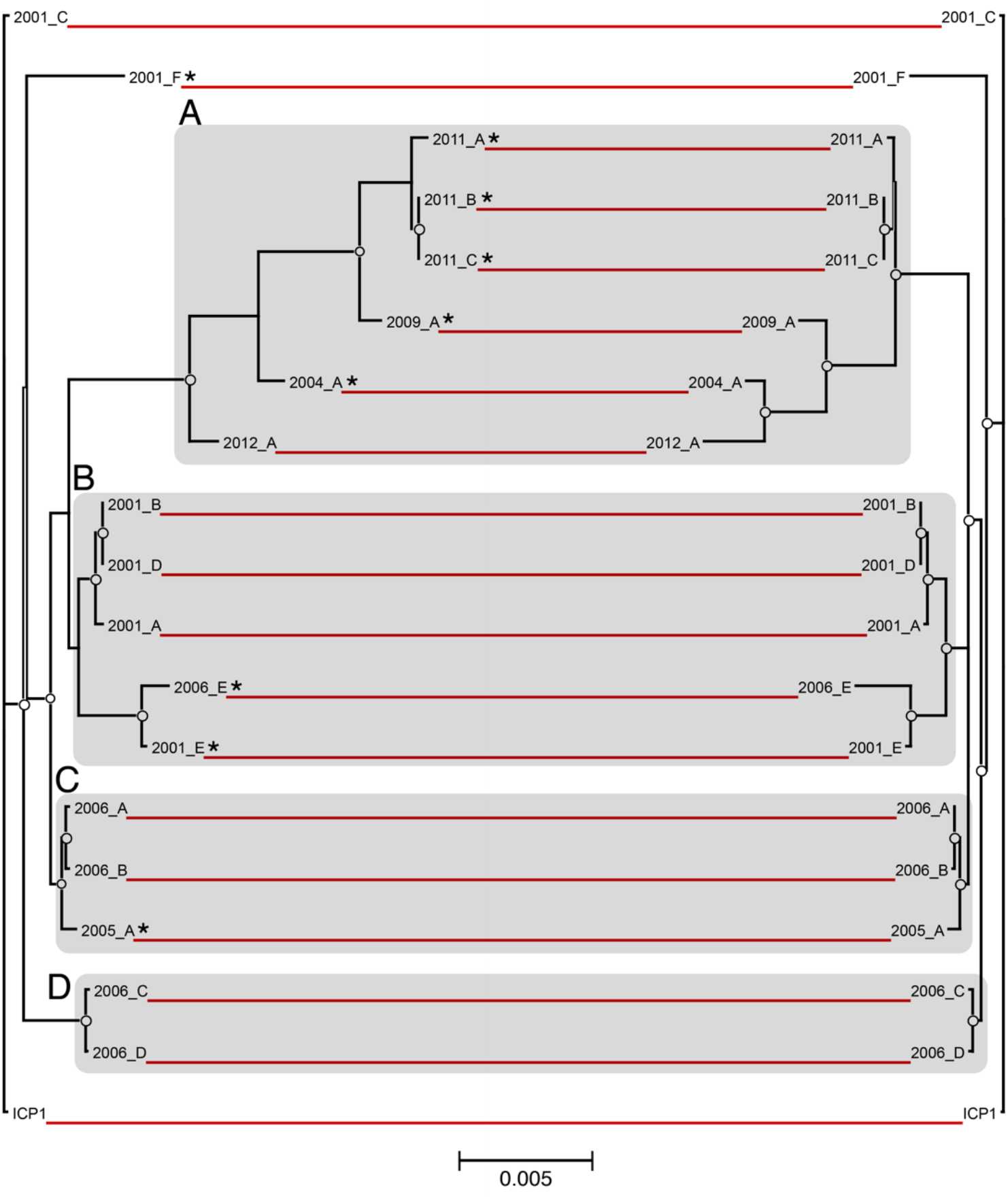
Phylogenetic comparison of ICP1 whole and core-genomes. Maximum likelihood phylogenetic trees (unrooted) were constructed based on a multiple whole-genome alignment of all 19 ICP1 phage sequences (left) as well as an alignment of the concatenated ORFs for each phage that comprise the core genome (right). The red lines connect identical leaves between trees to indicate relative phylogeny. The circles represent nodes with ≥90% bootstrap support (n=100). The scale bar measures nucleotide substitutions per base pair. Distinct phylogenetic clusters are shaded grey and CRISPR strains are marked with ‘*’.

Overall, the topologies between whole-genome and core-genome trees were highly similar with a few minor exceptions. Both share identical clustering and have almost no leaf-level differences within clusters B, C and D. Cluster A showed an inversion of the phylogenetic differences of the leaves between the two alignments. For instance, ICP1_2012_A has the most divergent core-genome (from all other strains) but is the least divergent within its cluster when the whole-genome was considered.

### 3.2. Genome Alignment Visualization

The consensus genomic sequence constructed from the whole-genome alignment was 151,832bp long and contained all of the coding and non-coding regions from each genome. It was used as a reference to map the genomes and visualize the overall multiple-genome alignment (Figure 2). Variable regions of insertions and deletions were visible as gaps in the circular alignment. Similar to the whole-genome phylogeny, there was not a clear progression of sequence divergence based on isolation chronology. No large regions of GC content difference were observed.

**Figure 2.**
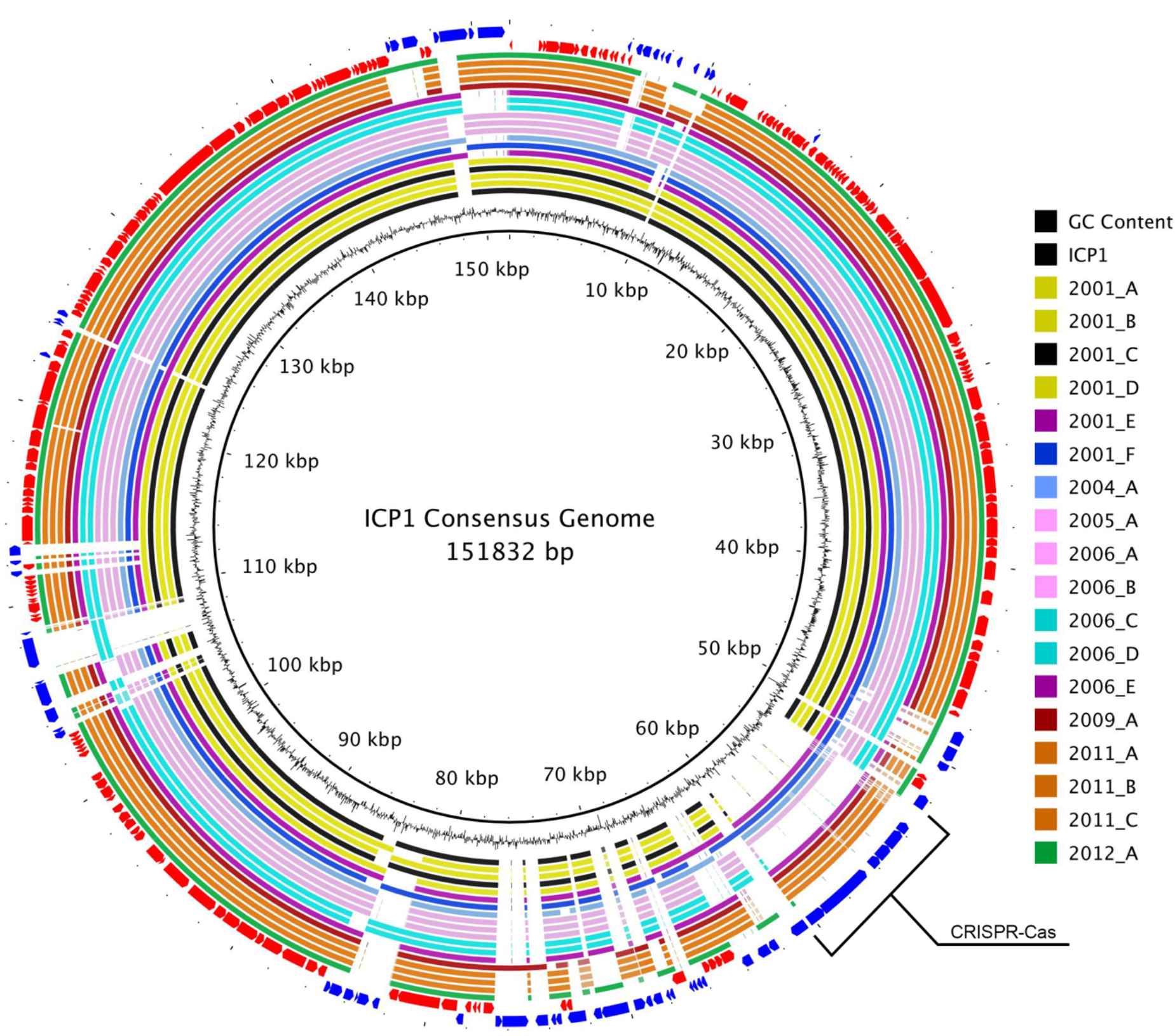
ICP1 pan-genome consensus alignment. A BLASTn-based whole genome alignment of all 19 ICP1 phage genomes using the MAUVE alignment consensus sequence as reference. The innermost ring is the consensus sequence. The next ring represents the GC content for that region. The following 19 rings display the alignment for each genome and are colored by phylogenetic similarity as determined by analysis of whole genomes (Figure 1 left). The second to last ring (red) represents the core-genomic ORFs while the outermost ring (blue) is the accessory-genome ORFs. The CRISPR-Cas insertion region is labeled.

### 3.3. ORF Annotation

Among all 19 genomes, a total of 269 distinct ORFs (based on homology cutoffs) were identified. Of these, 230 were originally annotated in the ICP1 ancestral genome and 39 were called by Prodigal. The number of ORFs per genome varied from 215 to 232 and demonstrated a slight but significant inverse relationship with year of isolation, i.e. more recent strains had fewer ORFs (Figure S1A). A weaker significant positive trend was observed between number of ORFs and genome length (Figure S1B) which is likely due to the CRISPR-Cas insertions, however there was not a statistically significant correlation between genome length and isolation year.

### 3.3. Core and Accessory Genome Analysis

To estimate gene diversity among the genomes we divided the total pan-genomic ORF complement into core and accessory groups. The core ORFs, those that were found in all 19 genomes, comprised ~70% of the total number of ORFs (185 out of 269) while the other 84 ORFs were considered to be accessory (Figure 3).

**Figure 3.**
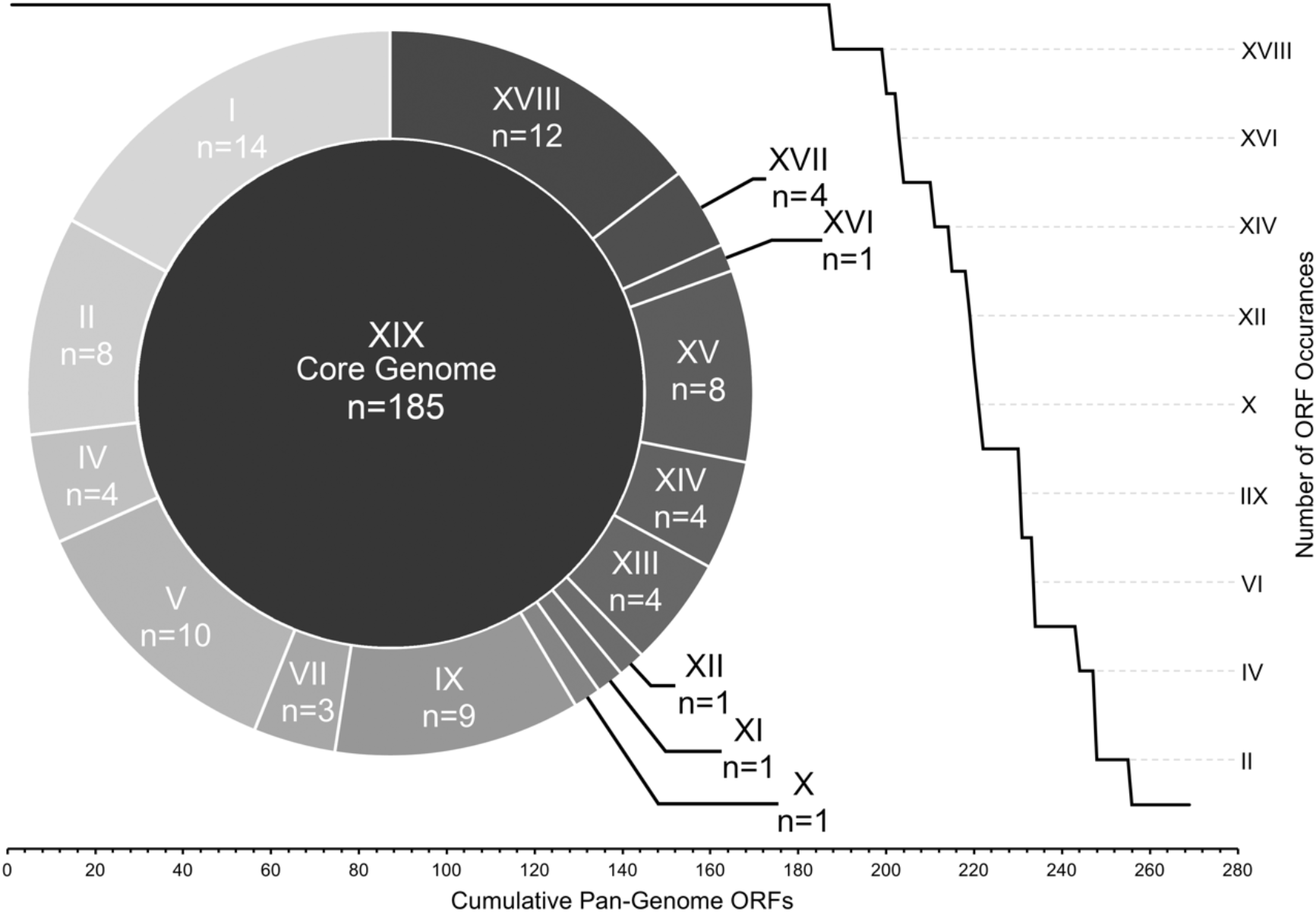
ICP1 pan-genome ORF allocation. All ICP1 ORFs arranged by number of genomes in which they were detected. ORF bins are represented by roman numerals, i.e. ‘XIX’ ORFs were found in all 19 genomes (core), ‘X’ ORFs were found in exactly 10 genomes, etc. The line graph demonstrates the overall cumulative curve of core and accessory ORF prevalence. The doughnut chart provides exact number of ORFs within each accessory bin.

Despite their occurrence across all genomes in this study, the core-genome ORFs were not all equally conserved at the sequence level. By comparing average pairwise nucleotide and amino acid sequence identity, we were able to resolve the core ORFs into three distinct groups: conserved-core, synonymous-core and divergent-core (Figure 4). The conserved-core was comprised of 49 ORFs that shared perfect sequence identity among all the genomic sequences. Correspondingly, they shared identical amino acid sequence identity. The synonymous-core included 26 ORFs that possessed a small amount of nucleotide diversity between genomes, but all mutations were silent, and the amino acid primary structure was therefore identical. Finally, the divergent-core contained the remaining 110 ORFs which were diverse in both nucleotide and amino acid pairwise identity. These divergent ORFs encompassed a range pairwise identity similarities from 99.97% to 91.57% at the nucleotide level and 99.98% to 91.83% for amino acid sequences (Table S1). The degree of pairwise identity difference was not correlated with an ORF’s sequence length (Figure 4).

**Figure 4.**
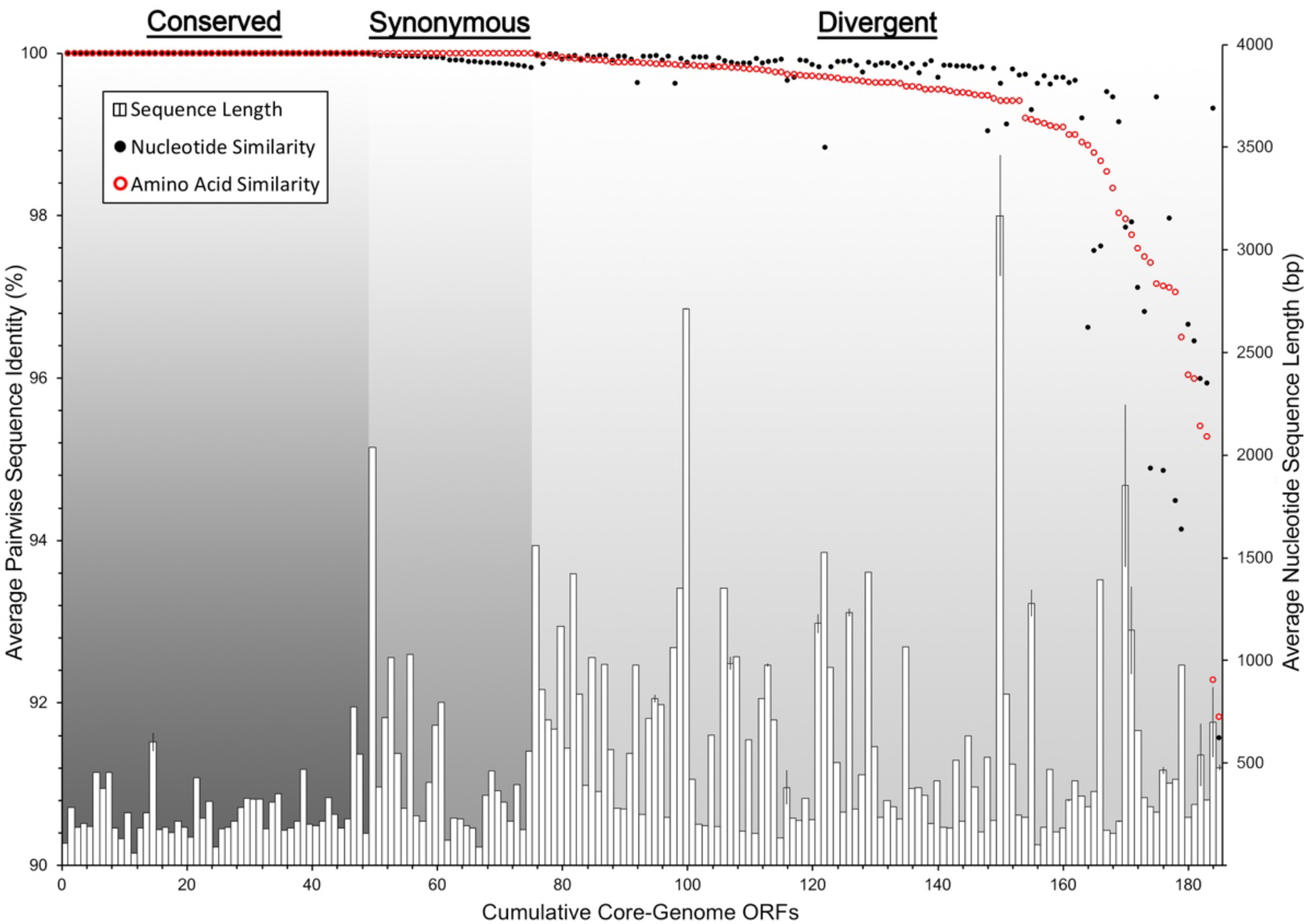
ICP1 core-genome ORF divergence. All ICP1 core-genome ORFs arranged by average pairwise similarity (171 pairwise comparisons) of both DNA nucleotide sequence alignments (black dots) and amino acid residue alignments (red circles). The histogram shows average nucleotide sequence length and standard deviation among the 19 sequences per ORF. The graph is divided into three sections by types of ORF similarity: Conserved (identical nucleotide and amino acid sequences), Synonymous (identical amino acid sequences, but silent nucleotide mutations) and Divergent (dissimilarity in both nucleotide and amino acid sequences).

The accessory ORFs were distributed among all 19 genomes in a complex manner (Table S2) and grouped into 15 levels of occurrence. These ranged from occurrences in 18 genomes (n=12) to singletons (n=14) (Figure 3) with several patterns of accessory ORF co-occurrence (Table S2). The most obvious example was the CRISPR-Cas locus which was previously known to reside in 8 of these genomes [13,15] and confirmed through our annotation methods. Other examples of sequential ORF co-occurrence included a locus containing ORFs 115, 116, 117, 117.1 and 118 which were found in the same five genomes, ORFs 160, 162, 163 in a different set of five overlapping genomes and ORFs 222, 223, 225 in 15 genomes.

### 3.3. Conserved Functional Domains

The majority of ORFs in both the core and accessory genomes were classified as hypothetical due to a lack of an informative BLAST identification. Only 18 of the core ORFs (9.7%) currently had a predicted function, two in the conserved-core, three in the synonymous-core and 13 in the divergent-core (Table S1). The accessory core contains 11 ORFs with a putative or known function (Table S2). The NCBI conserved domain search identified 18 additional core-genome ORFs and 11 accessory-genome ORFs that shared at least partial amino acid sequence homology to known functional domains.

## 4. Discussion

ICP1 is a *V. cholerae*-infecting bacteriophage that appears to be prevalent throughout the Bay of Bengal’s coastal areas and is transferred readily alongside *V. cholerae* into humans during cholera outbreaks in the region [11]. It is becoming increasingly theorized that this region is the source of most if not all global cholera outbreaks [3,4,27], and therefore a better understanding of this co-evolving predator is an important area of ongoing cholera research. In this study we have performed a comprehensive phylogenetic analysis on all available, well-sequenced ICP1 isolates to elucidate their genetic divergence over time and provide a platform on which to develop future ICP1-related bioinformatic analyses.

We have found that the genomes of ICP1 are surprisingly well-conserved between all isolates over the twelve-year period in which they were isolated. This is demonstrated in the whole-genome phylogeny which, while resolvable into distinct phylogenetic clusters, still only represents a maximum variation of approximately 1 nucleotide substitution per 100 base pairs between the most divergent isolates (Figure 1), many of which are likely silent or non-coding. A high degree of genomic conservation is also indicated by the relatively large core-genome shared between all isolates (Figure 3). This conservation is not only surprising due to the amount of time the core-genome has remained stable, but also due to the complex conditions that likely exist in the coastal and ocean environments where ICP1 is competing against a host population that is almost certainly more diverse than clonal outbreak strains. In at least one other study that examined multiple strains of a single marine bacteriophage, Far-T4, it was shown that sequence variability among strains was at least an order of magnitude greater than for ICP1 [28]. In contrast, a study from a less variable environment found that strains of a *Y. enterocolitica*-infecting podovirus were more highly conserved than ICP1 [29]. However, it must be considered that these environmental pressures may actually be what drives this bacteriophage to maintain a large core-genome, of which the vast majority of ORFs are hypothetical. This large gene complement may be providing the flexibility needed to compete in the Bay of Bengal. Currently we can only speculate on the exact reason for the conservation of ICP1’s genome, but expanding the repertoire of well-sequenced genomes will help to put a finer constraint on the core-genome and perhaps identify genetic targets for future study.

Despite the stable conservation of the core-genome, ICP1 also possesses a diverse collection of accessory genes that have been integrated into several locations around the pan-genome (Figure 2). These accessory ORFs appear to be both single acquisitions as well as integrations of larger loci (Table S2). The most notable of the latter is the previously described acquisition of a CRIPSR-Cas system which is used as a weapon in the arms race between ICP1 and *V. cholerae* [14,15]. This may suggest that other mechanisms of molecular warfare would be likely targets for acquisition, but if so, the conserved domain analyses failed to reveal mechanisms already known. Interestingly, though, the trend has been for genomic ORF counts to diminish over time culminating in the latest isolate from India in 2012 which possessed the fewest overall number of ORFs, i.e. the smallest accessory-genome (Figure S1). This could be due to shedding of outdated or detrimental genes, possibly because the acquisition of a CRISPR-Cas system provided enough flexibility to obviate the need for other mechanisms. And although ICP1_2012_A is CRISPR-Cas negative, the other five most recent isolates are positive. It is also possible that this is a regional difference though, and more sequenced genomes will help to determine if this trend is chronological or geographical or spurious.

Querying the ORFs against the NCBIs conserved domain database returned several hits having to do with replication, nucleotide metabolism, recombination, endonuclease activity and a few other basic functions (Table S1, S2). However, what may be most telling is that 80% of all ORFs do not possess homology to any conserved domain indicating that there is a great deal left to learn about the interaction between ICP1 and *V. cholerae*. It should also be noted that the annotation of ORFs necessarily requires certain assumptions to be made about similarity cutoffs and thresholds and as such these ORF calls are best viewed as estimates. However, we were conservative in our methods and are confident that the a very large proportion of the calls are accurate.

As we advance our understanding of how cholera spreads globally, it will be important to also continue tracking ICP1’s phylogeny and genetic composition so that we may develop a better understanding of its co-evolution with *V. cholerae* and attempt to disentangle the complex molecular and ecological interactions that may play an important role in defining cholera outbreaks.

## Supplementary Materials

The following are available online at www.mdpi.com/xxx/s1, Figure S1: ORF occurrence trends by year and genome length, Table S1: Core-genome ORFs pairwise similarity and CDD hits, Table S2: Accessory-genome ORFs ICP1 strain matrix.

## Author Contributions

Angus Angermeyer and Kimberley Seed conceived of the study; Angus Angermeyer analyzed the data; Moon Moon Das and Durg Vijai Singh contributed sequencing data and reagents. Angus Angermeyer and Kimberley Seed wrote the paper.

## Acknowledgments

We wish to thank Zach Barth for his helpful assistance and **Genotypic Technology, Bangalore, India** for the sequencing of ICP1_2012_A.

---

**Figure S1:**
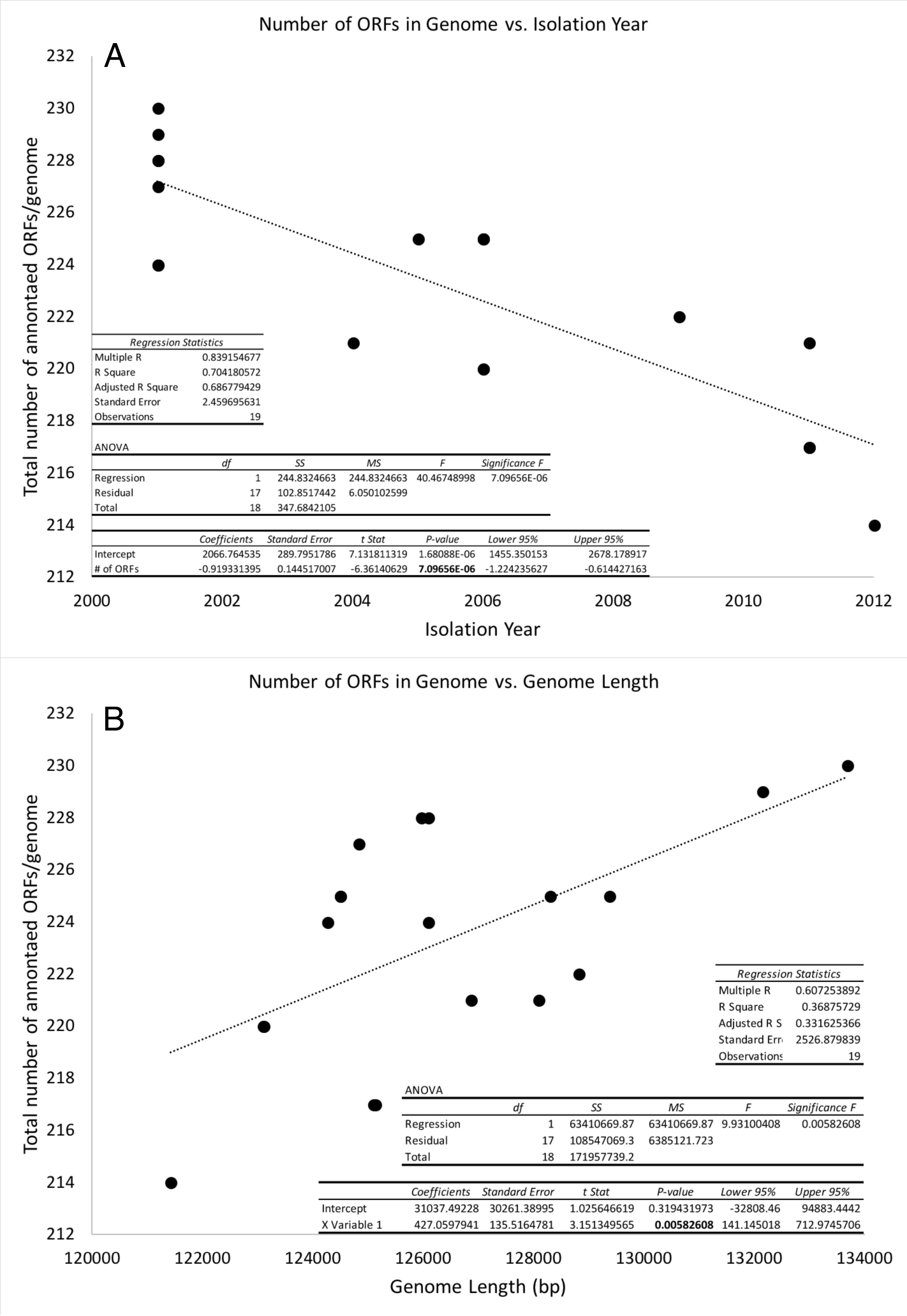
ORF linear regression statistics. ANOVA analyses of the linear regressions between (A) number of ORFs/genome vs genome isolation year and (B) ORFs/genome vs. genome length.

**Table S1:**
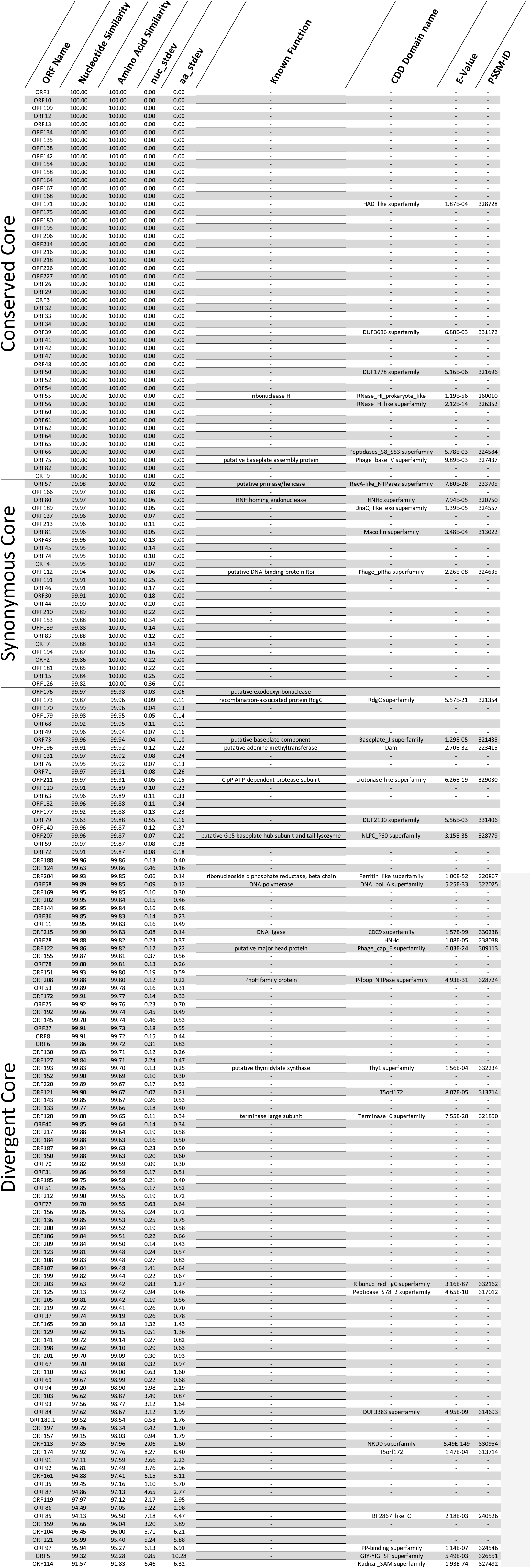
Core-genome information. The data for each ORF represented in Figure 4 is listed in columns 2 and 3. Columns 4 and 5 contain the standard deviations for those pairwise similarity values. The other columns contain information about putative gene function and conserved domain homology.

**Table S2:**
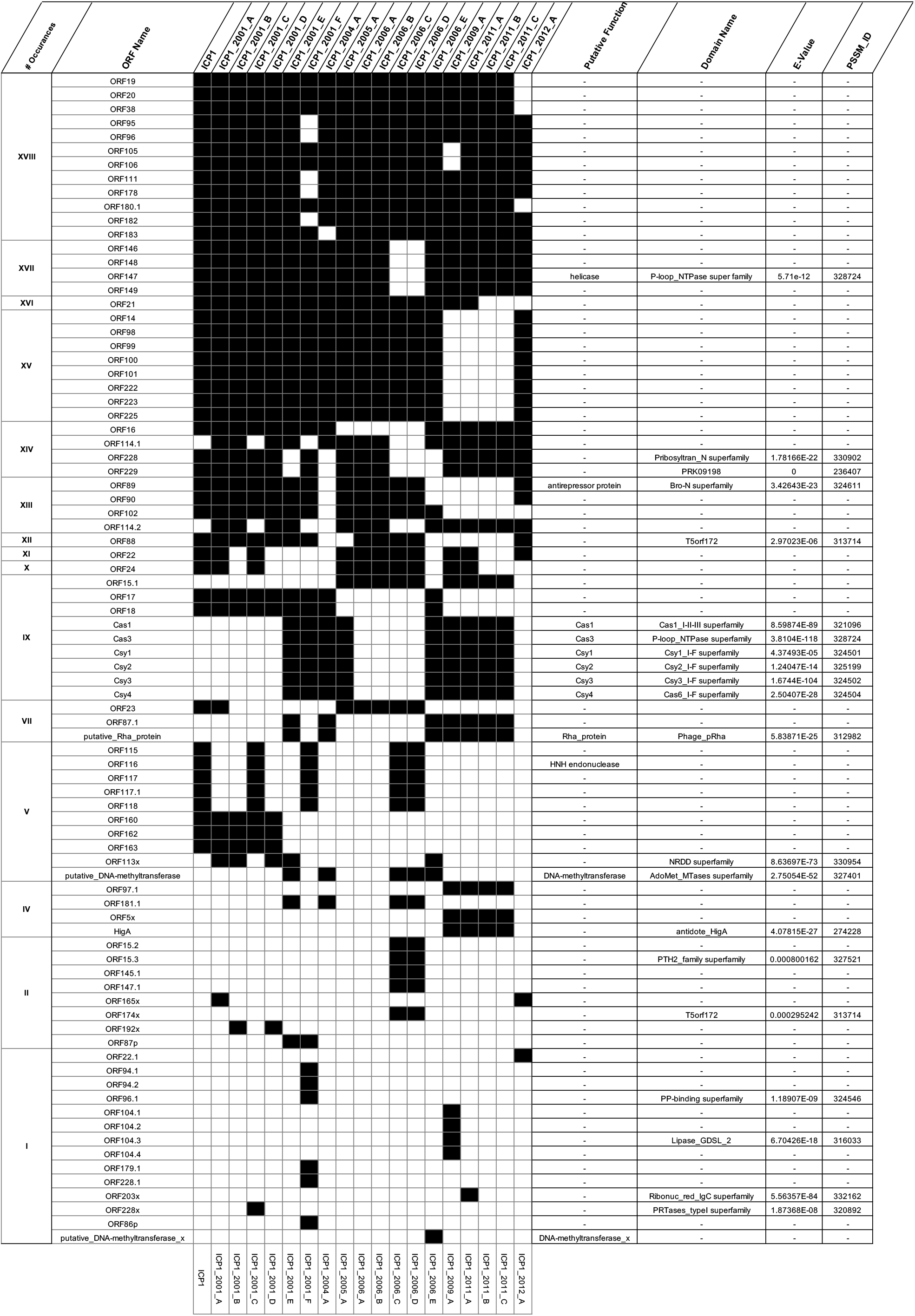
Accessory-genome ORF occurrence matrix. Each ORF in the accessory genome is listed along with any putative function or conserved domain homology information. A black square indicates that the ORF occurs in a specific genome.

